# Gene surgery as a potential treatment option for Nephropathic Cystinosis *in vitro*

**DOI:** 10.1101/2023.11.01.565117

**Authors:** E. Sendino Garví, J. Faria, C. Pou Casellas, S. Thijssen, E.J. Wubbolts, A. Jamalpoor, P. Harrison, R. Masereeuw, M.J. Janssen

## Abstract

Nephropathic cystinosis is a rare monogenetic kidney disease caused by mutations in the lysosomal transporter cystinosin (encoded by *CTNS*) that, to date, has no cure. The hallmark of this disease is lysosomal accumulation of cystine and decline in proximal tubular function leading to kidney failure early in life. In this project, we developed a novel gene repair strategy using CRISPR/Cas9 Homology-Independent Targeted Integration (HITI) to restore *CTNS*. A novel, non-viral peptide-mediated approach was used to deliver the Cas9-guideRNA ribonucleoprotein (RNP) complex and repair templates to conditionally immortalized proximal tubule epithelial cell (ciPTEC) lines. The repair constructs contained either mCherry (1.7 kb), the *CTNS* Superexon (1.7 Kb) or both (3.2 Kb). The results demonstrated that the smaller mCherry construct achieved a higher repair efficiency (63%) compared to the *CTNS*-mCherry construct (16%). Clonal expansion of repaired cells showed restoration of lysosomal cystine levels in 70-75% of the clones, which was accompanied by improved mitochondrial bioenergetics. In conclusion, CRISPR/Cas9 HITI can be used to precisely insert repair templates into the genome, resulting in a functional cystinosin restoration, and a reversal of the cystinotic disease phenotype.

## 1. Introduction

Cystinosis is a rare autosomal recessive genetic disorder that is characterized by the accumulation of cystine residues in the lysosomes, leading to progressive damage of various organs and tissues, including the kidneys [1]. Global estimates suggest that there are approximately 0.5–1 cases of cystinosis per 100,000 live births worldwide [2]. Nephropathic cystinosis, also known as infantile cystinosis, is the most frequent and most severe form and progresses into kidney failure and renal Fanconi syndrome [3], which eventually leads to the loss of barrier integrity of the proximal tubule starting at very young age [4–6], making early diagnosis and prompt treatment imperative for optimal patient outcomes. Cystinosis is caused by mutations in *CTNS* which encodes for the lysosomal membrane protein cystinosin and the plasma membrane protein cystinosin-LKG. Both isoforms function as a lysosomal cystine/proton symporter, and loss of *CTNS* results in lysosomal cystine accumulation and, eventually, organellar crystal formation [7,8]. Furthermore, cystinotic cells exhibit decreased ATP production and increased reactive oxygen species (ROS), aberrant mitochondrial function and size [9,10], in addition to increased apoptosis, altered mTORC1 activity, and impaired autophagy [11]. Although cystinosin-LKG has a targeting motive which initially directs it to the plasma membrane, studies have also found that it is able to move to the lysosomes though endocytic retrieval [12–14]. It is not known how the different isoforms contribute to the disease phenotype.

To date, a curative treatment for nephropathic cystinosis does not exist, however, the life expectancy of patients can be prolonged by the use of cysteamine and renal replacement therapy, in particular kidney transplantation [15]. Despite this improvement in quality of life, kidney transplantation is invasive and does not correct the systemic metabolic landscape of cystinosis, resulting in continuation of cystine accumulation in all cells of the body [16,17]. Cysteamine therapy is recognized as the first line of treatment for cystinosis by reducing the intralysosomal cystine content, thereby delaying the development of end stage kidney disease. However, the strict medicinal regimen and serious side-effects put a burden on the quality of life and, still, cannot prevent the loss of kidney function [17,18]. Although the kidney has inherent mechanisms that aim to restore its structure and function, the regenerative capacity of these cells is limited and often insufficient to maintain kidney function in severe conditions such as nephropathic cystinosis, where continuous and progressive damage occurs over time. The rapid development of CRISPR gene editing technologies offers a new path in which genetic (kidney) diseases could be targeted at its origin [19–21]. CRISPR technology is currently being evaluated in animals to treat a variety of genetic diseases, including Huntington’s disease and muscular dystrophy [22–24]. Additionally, this revolutionary technology has been successfully tested in humans to treat β-thalassemia, sickle cell disease [25] and HIV-1 [26]. Since the discovery of CRISPR gene editing tools, many variations and applications have been designed to achieve more specific edits, enhance efficiencies, increase specificity, and minimize off-target effects [27]. Homology-independent targeted integration (HITI) is a CRISPR/Cas9 technique that enables targeted integration of exogenous DNA into a specific genomic locus without relying on homology-directed repair (HDR). This technique offers several advantages over HDR-based methods, such as a higher efficiency of integration, a reduced risk of off-target effects and the ability to edit both dividing and non-dividing cells [28]. Moreover, CRISPR HITI could be used to insert entire genes into a specific site in the genome, indicating its potential for gene therapy applications [28–30].

The most common mutation in cystinosis patients of Northern European ancestry (in over 50% of mutant alleles) is a 57-kb deletion that completely removes the first 10 exons of *CTNS* [31]. CRISPR HITI could be used to precisely integrate the first 10 exons of the *CTNS* gene, thereby restoring gene function for any mutation located in those regions. The advantage of this approach is that the natural splicing of exon 11 and 12 will be maintained and the cells will be able to produce both functional isoforms of cystinosin. This novel technology, combined with the efforts in engineering tolerable and efficient delivery methods, could open doors for CRISPR-based therapies to enter the clinics [32].

In this study, we aimed to test a potential curative therapy for nephropathic cystinosis *in vitro* using CRISPR/Cas9 HITI technology in conditionally immortalized proximal tubule epithelial cells (ciPTEC). We designed and generated several DNA repair templates for the *CTNS* gene and introduced these repair cassettes into cells using a cell penetrating peptide. We show that the repair of the *CTNS* gene using CRISPR/Cas9 HITI can restore the gene function and reduce lysosomal cystine accumulation *in vitro*.

## 2. Materials and Methods

### 2.1 Cell culture of ciPTEC

Three different ciPTEC were used in this study. The ciPTEC *CTNS*^-/-^line was previously generated in our group from the healthy ciPTEC 14.4 (ciPTEC *CTNSWT*) as described by Jamalpoor et al. [11]while the ciPTEC *CTNS^∆57kb^* line was generated from a cystinotic patient as described by Wilmer et al. [33]. All ciPTEC lines were cultured according to the method previously described [33]. In brief, Dulbecco’s modified Eagle medium (DMEM/F-12; GIBCO), supplemented with 10% (v/v) fetal calf serum, 5 μg/ml insulin, 5 μg/ml transferrin, 5 μg/ml selenium, 35 ng/ml hydrocortisone, 10 ng/ml epidermal growth factor, and 40 pg/ml triiodothyronine, was used as the culture medium. Cells were seeded at a density of 48,400 cells/cm2 and initially grown at 33 °C for 24 h to promote proliferation, followed by incubation at 37 °C for 7 days to allow for full differentiation into epithelial cells.

### Repair constructs and transfections

Both the 1.7Kb mCherry and the 1.7Kb Superexon HITI templates were purchased from IDT (Integrated DNA Technologies BV). Since the 3.2Kb mCherry_Superexon HITI template could not be provided by an external supplier, this template was generated by PCR using the Platinum Super-FiII high-fidelity enzyme (Thermofisher Scientific, #<u>12361010</u>) from a plasmid purchased from Genscript (Genscript Biotech Corporation) using the following primer set: 5’-GAATTCGAGCTCGGTACCTC-3’ as forward primer and 5’-CTTGCATGCAGGCCTCT-3’ as reverse primer. As a control to account for possible effects of the different origin of the HITI repair templates (in-house PCR-based versus commercially manufactured), a 1.7Kb mCherry template was generated by high-fidelity PCR from the commercial template using 5’-GAATTCGAGCTCGGTACCTCC-3’ and 5’-AGAGGCCTGCATGCAAG-3’ as forward and reverse primers, respectively. The custom gRNA (5’-CCCAGTCAGAACGCTGGCCCTCC-3’) designed to target the intronic region between exons 10 and 11 of the *CTNS* gene were ordered from IDT (Integrated DNA Technologies BV) and SpCas9 was ordered from Sigma (Sigma Aldrich, #CAS9PL). Prior to transfection, ciPTEC *CTNS*^-/-^ and ciPTEC *CTNS^∆57kb^* cells were seeded in 96 well plates at a density of 48,400 cells/cm2. After 24h at 33 °C, SpCas9 (final concentration of 50μM) and gRNA (final concentration of 50μM) were mixed and incubated for 10 min at room temperature. Afterwards, LAH5 peptide (BIOMATIK) was added to a final concentration of 5μM and incubated for 15 more min at room temperature. Immediately after this incubation, 20μg of the corresponding HITI repair templates, followed by Optimem media (GIBCO), were added and this mixture was added to the wells. After 48 h cells, were collected for clonal expansion or flow cytometric analysis.

### Fluorescence-activated cell sorting (FACS) and flow cytometry

For sub-clone generation, cells were harvested with Accutase (Thermofisher Scientific) and resuspended in their normal culture media 48 h after transfection. Cells were centrifuged at 300xg for 3 minutes and resuspended in their normal culture media. The cell suspension was then put through a 70 microns cell strainer (Corning) and they were sorted into single cells by sorting one cell per well in a 96 well plate (Greiner) with the BD FACSAria™ III Cell Sorter (BD Biosciences) using the BD FACSDiva™Software. For cells transfected with mCherry_Superexon, the mCherry positive cells were selected for clonal expansion and, for the cells transfected with the *CTNS* Superexon, cells were randomly selected for clonal expansion. In total, 96 cells were sorted in each condition and all the clones that survived the cloning process (38 in total) were expanded for future experiments.

To evaluate integration efficiency after transfection, mCherry positive cells were quantified using flow cytometry. Cells were processed as mentioned above until the step before putting them through the cell strainer, which was done with PBS instead of culture media, after which the cells were centrifuged at 300xg for 3 minutes and subsequently resuspended in 4%PFA (Pierce™ 16% formaldehyde (w/v), methanol-free, Thermofisher Scientific) for 20 min at room temperature. After incubation, cells were centrifuged 3 min at 300xg and resuspended in PBS with 1% BSA (Sigma Aldrich) and 1mM EDTA (Gibco) for analysis using the 610/20 filter for the Yellow/Green laser (561 nm) on the CytoFLEX LX (Beckman Coulter, Brea, CA, USA). The flow cytometry data was analyzed using the software (FlowLogic Alias version 8.1, INIVAI technologies, Australia).

### Estimation of HITI efficiency by PCR

To investigate in which direction the HITI templates were inserted into the genome of the transfected cells and to investigate mono-allelic and bi-allelic insertion, the DNA of the cells was collected using QIAamp DNA Mini Kit (QIAGEN, #51304). The *Q5* DNA polymerase (New England Biolabs) was used for region amplification using specific primes (Table 1) by PCR. Amplicons were purified using QIAquick PCR purification kit (QIAGEN, #28104). The samples were then mixed with Gel Loading Dye, Purple (6X) (New England Biolabs) and loaded in a 2% Agarose gel containing SYBR Safe DNA gel stain (Thermofisher Scientific, #S33112) and ran for 45 min electrophoresis at 120V. For DNA band visualization, the gels were transferred to the GelDoc Go System(Biorad) and imaged following the pre-set for SYBR Safe-stained gels.

**Table 1.**
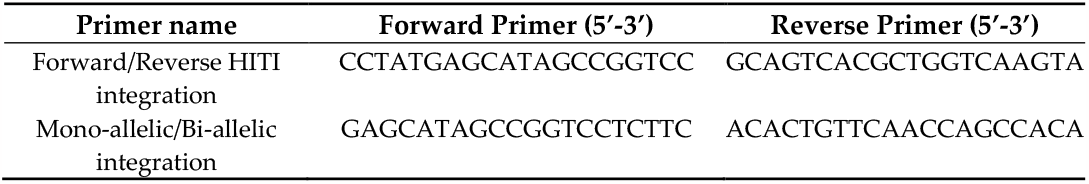
Primer sequences for HITI efficiency detection.

**Table 2.**
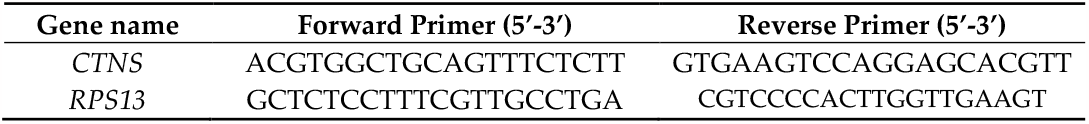
Primer sequences for Real-time qPCR.

### Quantification of mRNA expression with Real-time qPCR

The mRNA was extracted from the cells using the Qiagen RNeasy mini kit (QIAGEN), followed by reverse transcription of a total of 600ng of mRNA using iScript Reverse Transcriptase Supermix (Biorad). The expression levels of the *RPS13* gene were used as normalization housekeeping gene within the same sample (∆Ct), and the fold-change in expression levels were calculated using the 2-ΔΔCt method. The mRNA expression levels of *CTNS* and *RPS13* were determined using quantitative real-time PCR with iQ Universal SYBR Green Supermix (Biorad). For *CTNS* quantification, primers were designed to span exon 10 and 11 that will only amplify if the complete mRNA transcript is found.

### Quantification of lysosomal cystine using LCMS

Cystine levels were measured using an in-house developed and validated HPLC-MS/MS assay by Jamalpoor et al. (2018) [34]. The ciPTEC were seeded in 6 wells plates (Greiner) and transferred to 37 °C after 24h at 33 °C. Cell pellet was collected and suspended in N-Ethylmaleimide (NEM) solution (Sigma Aldrich), and the resulting cell suspension was precipitated, and protein was extracted using sulfosalicylic acid 15% (w/v). Protein concentration was determined using the PierceTM BCA protein assay kit (Thermofisher), and cystine concentration was measured using HPLC-MS/MS. The data is presented as cystine values (nmol) normalized to the total protein content of each sample.

### Mitochondrial respiration assessment with Seahorse assay

To perform the Seahorse assay, we first seeded 2,000 cells in specialized Seahorse assay plates (Seahorse XF Cell Mito Stress Test Kit, Agilent) and incubated them for 24h at 33 °C for proliferation and for 7 days at 37 °C for maturation. On the day of the assay, cells were washed with Seahorse assay medium, and loaded into the plate with appropriate amounts of provided compounds or inhibitors according to the manufacturer’s protocol. The assay plate was then placed in the Seahorse instrument (Agilent Seahorse XF Pro) to measure the oxygen consumption rate (OCR) and extracellular acidification rate (ECAR) of the cells in real-time. We analyzed the Seahorse data using GraphPad Prism version 9.3.1 (GraphPad Software, Inc., USA). In brief, the baseline respiration is measured whereafter oligomycin is added, to access the ATP production. Subsequently, carbonyl cyanide-p-trifluoromethoxyphenylhydrazone (FCCP), rotenone and antimycin, provided with the assay kit, are administered to determine the proton leak and the levels of maximal and spare respiration capacity. Furthermore, a high OCR and low ECAR are indicative of cells that are highly active and efficient at producing energy aerobically, while a low OCR and high ECAR indicate that cells are primarily relying on anaerobic metabolism and may be under stress or facing energy limitations.

### Gaphs and statistical analysis

All graphical illustrations and statistical analyses were performed using GraphPad Prism version 9.3.1 (GraphPad Software, Inc., USA) unless otherwise specified. For graphical presentation, the data is shown as the mean ± standard error of the mean (SEM), including the individual values as dots. All experiments were performed in three independent experiments and in technical duplicates, unless specified otherwise. Outliers were excluded based on their distance to the median. Statistical significance was evaluated using One-way Analysis of Variance (ANOVA) or two-tailed Student’s t-test, when appropriate. A p-value of < 0.05 was considered significant.

## 3. Results

### Fluorescent detection of CRISPR/Cas9-HITI insertion efficiency

We designed a *CTNS* repair construct (Superexon, 1.7 Kb) containing the endogenous *CTNS* promoter, the first 10 exons of the *CTNS* gene, and the splicing donor site of exon 10, resulting in a linear double stranded DNA sequence without bacterial or other exogenous DNA sequences. Inserting this repair construct in the intronic region before exon 11 should allow for dynamic expression regulation and alternative splicing of exon 11 and 12, depending on the requirements of the cell (Figure 1A). To optimize the delivery of this construct in target cells, we also created two constructs to allow for fluorescent detection of successful target integration: one mCherry construct with the same size as the Superexon (mCherry, 1.7 Kb) and one construct with both mCherry and the Superexon construct (mCherry_Superexon, 3.2 Kb) (Figure 1B). In line with the HITI strategy used to design our repair sequences [29], the gRNA sequences flanking the constructs were placed in the reverse direction (relative to the genomic orientation) at both ends of our repair templates to facilitate correct orientation of our repair template (Figure 1B-C).

**Figure 1.**
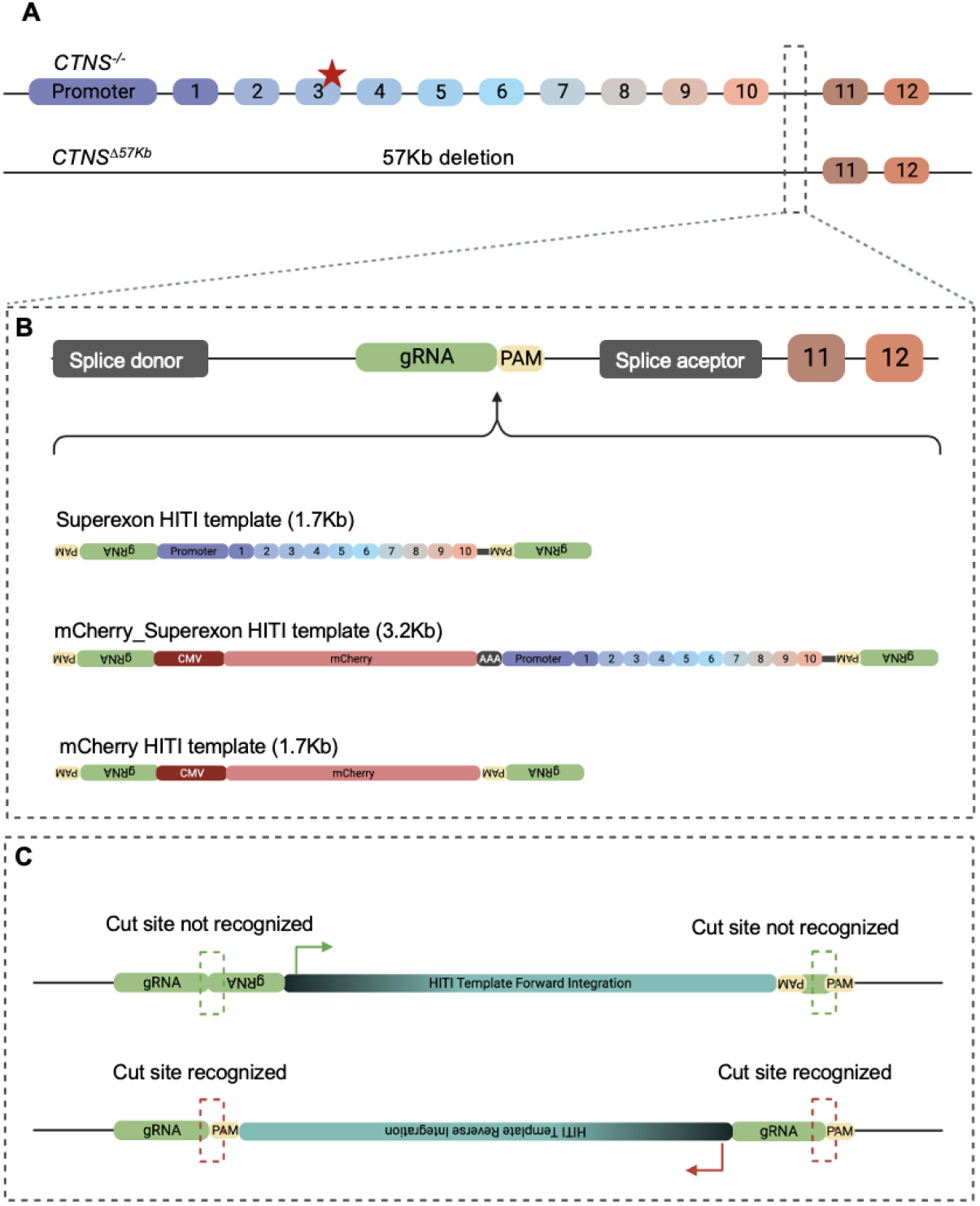
Schematic representation of the *CTNS* CRISPR/Cas9 HITI repair strategy. (**A**) Schematic overview of the *CTNS* gene sequence in the ciPTEC *CTNS*^-/-^ and ciPTEC *CTNS^∆57kb^*. CRISPR/Cas9 HITI was designed to target the genomic region between splicing sites in between exons 10 and 11 (dashed box). Red star indicates the location of the mutation present in ciPTEC *CTNS*^-/-^. (**B**) Schematic representation of the three HITI templates designed for this study: Superexon, mCherry_Superexon, and mCherry. (**C**) Schematic representation of the insertion directionality of the repair sequences and HITI. After Cas9 cuts both the HITI templates and the genomic region between exons 10 and 11, the HITI templates can get inserted in either direction. Because of the HITI design, if the templates get inserted in the forward direction, the insertion region will not get recognized by Cas9/gRNA and will stay inserted in the forward direction. On the contrary, if the repair templates get inserted in the reverse direction, two new cut sites will become available and Cas9 will cut again, strongly favoring integration in the forward direction.

Next, we investigated the insertion efficiency of the repair templates in three different proximal tubular cell lines: a control ciPTEC line with normal *CTNS* function (ciPTEC *CTNS*WT), a *CTNS* knockout generated from ciPTEC *CTNS*WT using CRISPR/Cas9 (ciPTEC *CTNS*^-/-^) and a ciPTEC line generated from a cystinosis patient carrying a 57Kb deletion (ciPTEC *CTNS^∆57kb^*) (Figure 2). As the 3.2Kb mCherry_Superexon HITI template could not be commercially manufactured due to its size, we purified its sequence in-house from a custom plasmid by high-fidelity PCR (Figure 2A). The 3.2Kb mCherry_Superexon HITI template achieved 12-15% insertion efficiency (Figure 2B) compared to a 63% and 48% insertion efficiency for the 1.7Kb mCherry PCR-based HITI sequence in the ciPTEC *CTNS*^-/-^ and ciPTEC *CTNS^∆57kb^*, respectively (Figure 2C-D). This indicates that the smaller sequence is more efficiently inserted into the host genome.

**Figure 2.**
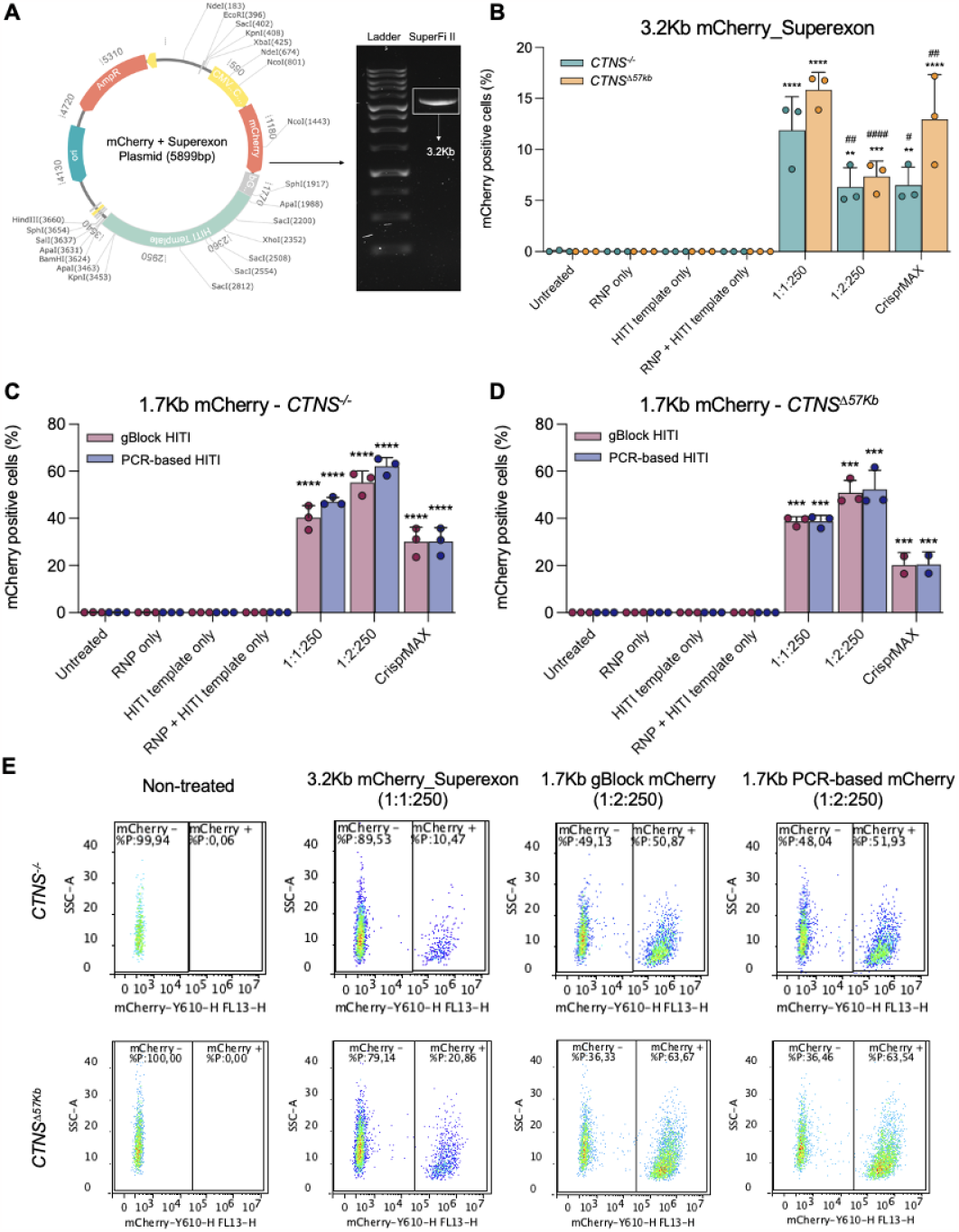
The *CTNS* CRISPR/Cas9 HITI templates achieve up to 60% of insertion efficiency in cystinotic ciPTECs. (**A**) Successful extraction and purification of the 3.2Kb mCherry_Superexon HITI repair template from the original plasmid by high-fidelity PCR. (**B**) Insertion efficiencies of mCherry based constructs evaluated by flow cytometry for the 3.2Kb mCherry_Superexon HITI template (PCR-based). The insertion efficiencies of the 1.7Kb mCherry HITI template (PCR-based) and the 1.7Kb mCherry HITI template (gBlock) were evaluated in ciPTEC *CTNS*^-/-^ (**C**) and ciPTEC *CTNS^∆57kb^* (**D**). For the peptide delivery different amounts of HITI template were used as indicated by the 1:1:250 and 1:2:250 which correspond to the different molar ratios the individual components: RNP:HITI template:LAH5 peptide. Delivery with CrisprMax was included for comparison. All conditions we tested in both ciPTEC *CTNS*^-/-^ and ciPTEC *CTNS^∆57kb^*. Flow cytometry experiments have been performed in triplicate with 2 technical replicates, representative flow cytometry plots are shown in panel **E**. (One-way ANOVA, *= p-value<0.05, **/## = p-value<0.01, *** = p-value<0.001, **** = p-value<0.0001). “*” represent comparisons between the untreated condition and the other conditions, “#” represent comparisons between ciPTEC *CTNS*^-/-^ and ciPTEC *CTNS^∆57kb^* amongst the different conditions.

For gene editing applications, the use of gBlock is preferred as this can provide a flexible high-quality sequence verified source of double stranded DNA. To exclude any variation in the production of the plasmid, we compared the efficiency for the PCR-based with the gBlock (purchased) HITI templates for the 1.7Kb mCherry template (Figure 2C-D). Insertion efficiencies of the gBlock template reached 58% and 46% in ciPTEC *CTNS*^-/-^ and ciPTEC *CTNS^∆57kb^*, respectively; and 63% and 48% in the case of the PCR-based template (Figure 2C-D). Our results showed no differences in the insertion efficiencies between gBlock and PCR-based templates, and therefore any further experiments using the smaller fragments (both mCherry and Superexon) were performed using gBlock templates.

### High percentage of functional repair after CRISPR HITI with the CTNS Superexon

To investigate the efficiency of targeted integration and restoration of the *CTNS* function using CRISPR HITI, cells were sorted 48 h after transfection into single cells for clonal expansion. In the absence of a fluorescent marker gene, the Superexon treated cells were randomly selected, whereas for the mCherry_Superexon the mCherry positive cells were selected (Figure 3). After expansion, we assessed the restoration at a functional level, including the *CTNS* mRNA expression levels and cystine accumulation of the individual clones (Figure 3A-D). The primer design to detect the mRNA expression of the *CTNS* gene also allowed us to detect the integration of the repair construct at its 3’ end, since the sequence amplified from these primers spans the nucleotides between the 3’ end of exon 10 and the 5’ end of exon 11. This means that the insert will only be amplified if it integrates into the correct spot in the genome and gets sliced to form the mature *CTNS* mRNA. In ciPTEC *CTNS*^-/-^, mRNA was still produced by the mutated allele (Figure 3A-B), while the ciPTEC *CTNS^∆57kb^* mRNA expression was detected only after insertion of the repair construct (Figure 3A-B), which correlated well with a reduction in cystine levels (Figure 3C-D), confirming functional restoration of the cystinosin transporter. Although *CTNS* gene expression did not reach the level of the healthy control, an increase of 5-10% in mRNA seemed sufficient to restore cystine levels around or below the plasma levels found in healthy individuals (Figure 3C-D). Lastly, we observed the reduction in lysosomal cystine accumulation in 70% of the randomly, blinded, sorted clones and in 75% of the mCherry-positive clones, when compared to the healthy ciPTEC *CTNS*WT line (Figure 3C-D), confirming the much higher insertion efficiencies of the smaler Superexon HITI template.

**Figure 3.**
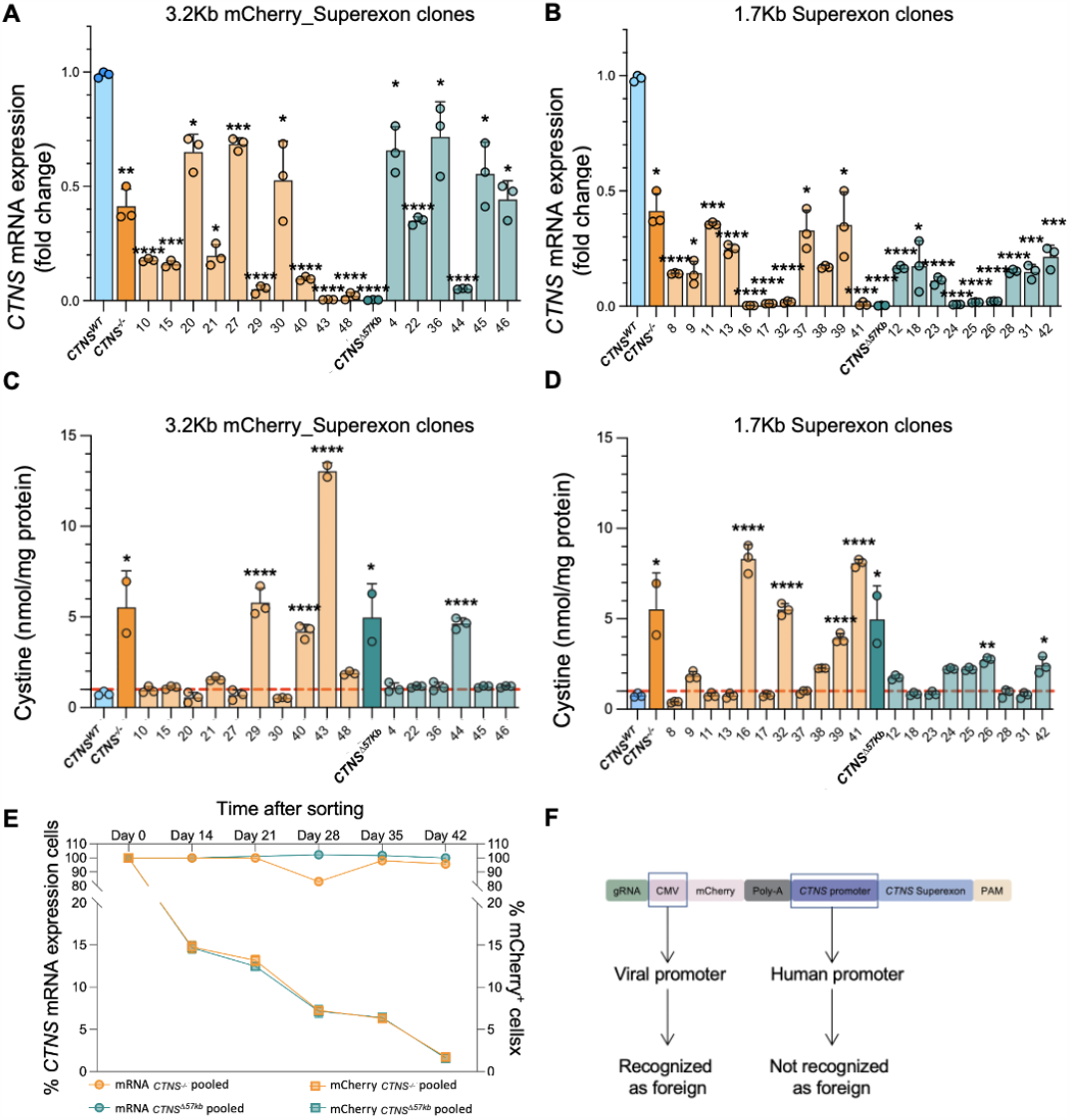
*CTNS* gene repair improved mRNA expression levels and cystine accumulation in cystinotic. The mRNA expression of the *CTNS* gene in mCherry-sorted clones transfected with the 3.2Kb mCherry_Superexon HITI template (**A**) or blind sorted clones after transfection with the 1.7Kb Superexon HITI template (**B**). Cystine levels after transfection with the mCherry_Superexon HITI template (**C**) or the Superexon HITI template (**D**). A total of 20000 cells were sorted, from both ciPTEC *CTNS*^-/-^ and ciPTEC *CTNS^∆57kb^*, for mCherry expression 48 h after transfection. The mCherry expression decreased over time while the mRNA expression of the *CTNS* gene within the same construct remained constant (this experiment was performed once) (**E**). Schematic simplified representation of the 3.2Kb mCherry_Superexon HITI template: the mCherry gene is expressed via the viral CMV promoter while the *CTNS* gene in the same construct is expressed via the *CTNS* human promoter (**F**). The red dotted horizontal line represents the cystine values measured in plasma of healthy individuals (<1nmol/mg) [18]. (* = p-value<0.05, ** = p-value<0.01, *** = p-value<0.001, **** = p-value<0.0001).

Furthermore, we designed specific primers to investigate the incidence in which the HITI construct was inserted in the reverse orientation or only in one allele (Figure A1). Out of the 16 mCherry sorted clones (transfected with 3.2Kb mCherry_Superexon HITI), only one (clone 48) integrated the repair sequence in the reverse direction (Figure A1A). Additionally, we designed primers that cover the breakpoint before integration to detect the absence of any integrations (Figure A1B). For all clones (except for clone 29) a band was detected at the expected size, suggesting that at least one allele was unchanged and in the case that a repair took place, it was in only one allele.

After integration of the 3.2Kb mCherry_Superexon HITI template we observed that over time the mCherry signal reduced from 100% after sorting to less than 5% within 42 days. As the *CTNS* Superexon and mCherry are located on the same construct, we evaluated whether the *CTNS* levels would also change over time as well. Gene expression analysis showed stable expression of *CTNS* mRNA, suggesting that, in contrast to the endogenous *CTNS* promotor, the CMV promotor was prone to get silenced (Figure 3E-F).

### Restoration of CTNS function improves mitochondrial respiration landscape

Cystinosis is a metabolic disorder in which the mitochondria are crucial in redox imbalances initiated by cystinosin dysfunction. To further assess the repair in the cystinotic models, we evaluated the mitochondrial function, including the respiration capacity, the proton leak and glycolysis. For this, we selected four clones of each cystinotic model, two that previously showed to be not corrected and two that showed to be corrected, based on their *CTNS* mRNA levels and lysosomal cystine values. For clones that originated from ciPTEC *CTNS*^-/-^, clones 10 and 8 were considered corrected and clones 43 and 16 were considered not corrected. Similarly, clones 45 and 18 were considered corrected and clones 44 and 24 were considered not corrected from the ciPTEC *CTNS^∆57kb^* derived clones.

The Seahorse assay determines the oxygen consumption rate (OCR) and extracellular acidification rate (ECAR) of the cells. The OCR is a measure of the rate at which cells consume oxygen in the process of cellular respiration, while the ECAR refers to glycolysis. We used these measurements to determine the fitness of the cells in terms of their energy production and usage. Figures 4A and B show an overview of the bioenergetic state of the ciPTEC *CTNS*^-/-^ derived clones and ciPTEC *CTNS^∆57kb^* derived clones, respectively. Increased OCR values in the corrected ciPTEC *CTNS^∆57kb^* derived clones (45 and 18) was observed when compared to the uncorrected clones and the ciPTEC *CTNS^∆57kb^* line (Figure 4B). While the maximal respiration capacity and proton leakage of the mitochondria was comparable amongst corrected and uncorrected ciPTEC *CTNS* -/- clones, and the ciPTEC *CTNS*^-/-^ line (Figure 4C-D), the corrected ciPTEC *CTNS^∆57kb^* derived clones showed an increase in maximal respiration capacity and a significant decrease in proton leak, indicating an improvement of the mitochondrial respiratory conditions, when compared to the ciPTEC *CTNS^∆57kb^* line (Figure 4E-F).

**Figure 4.**
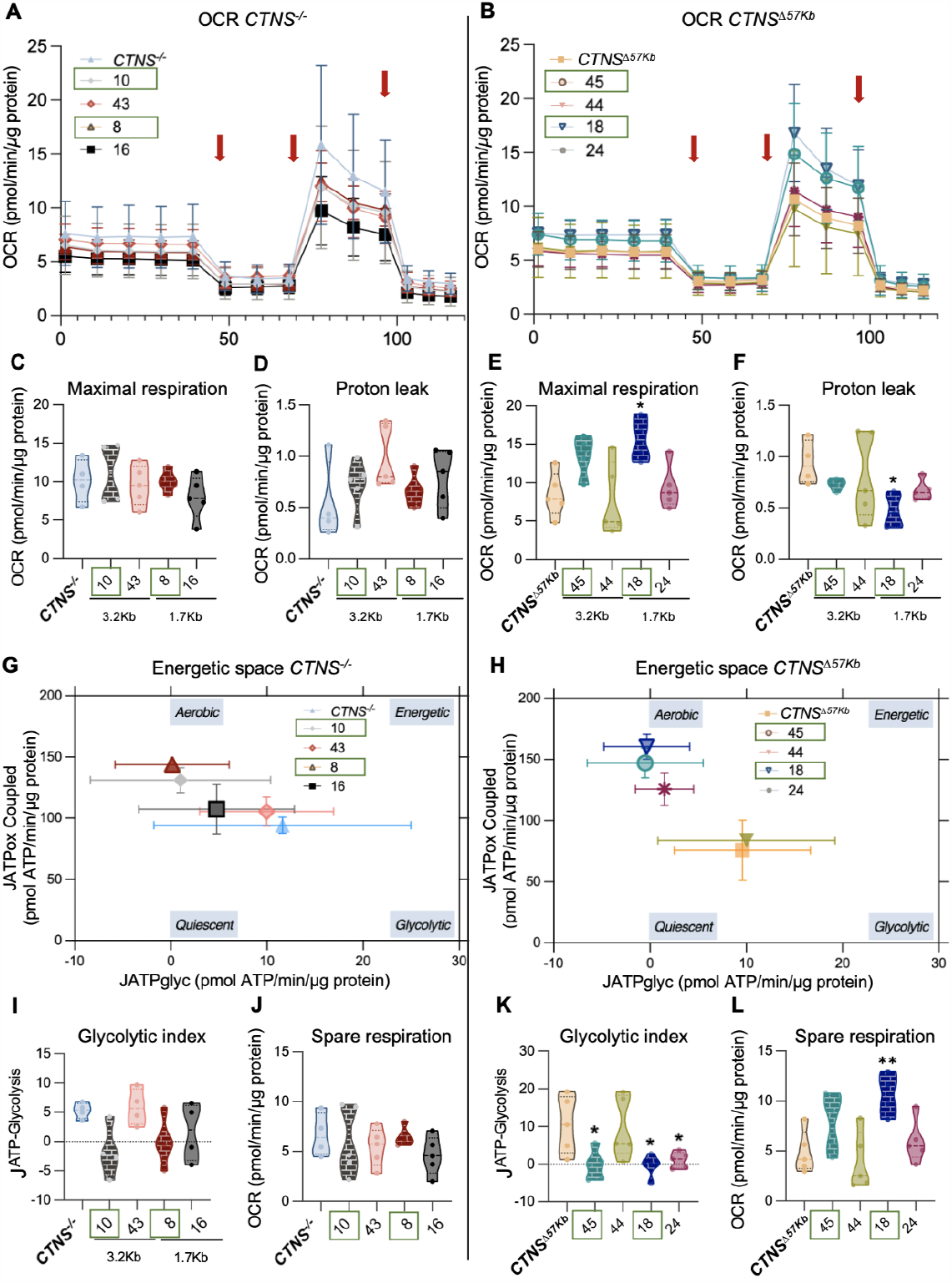
Effect of functional *CTNS* restoration on mitochondrial respiratory fitness and bioenergetic space. (**A-B**) Overview of the oxygen consumption rate (OCR) of the selected clones after stimulation with oligomycin, FCCP, rotenone + antimycin at the time points in the graphs indicated with red arrows, in the mentioned order. From the OCR overview graph, the maximal respiration capacity and the proton leak can be calculated. (**C-F**) Maximal respiration and proton leak of ciPTEC *CTNS*^-/-^ and ciPTEC *CTNS^∆57kb^* derived clones, respectively. (**G-H**) Energetic space of ciPTEC *CTNS*^-/-^ and ciPTEC *CTNS^∆57kb^* derived clones, respectively. From the Energetic space overview graph, the glycolytic index and the spare respiration can be calculated. (**I-L**) Glycolytic index and spare respiration of ciPTEC *CTNS*^-/-^ and ciPTEC *CTNS^∆57kb^* derived clones, respectively. Green box indicates “corrected clones” based on previous mRNA and cystine values (* = p-value<0.05, ** = p-value<0.01).

Net rates of ATP production derived from oxidative reactions and glycolysis were calculated from OCR and ECAR to allow visualization of the bioenergetic space of each clone (Figure 4G-H). We observed that the corrected ciPTEC *CTNS*^-/-^ derived clones (10 and 8) were located towards a more aerobic energetic space when compared to the uncorrected clones and the ciPTEC *CTNS*^-/-^ line (Figure 4G). Similarly, the corrected ciPTEC *CTNS^∆57kb^* derived clones (45 and 18) showed to be in an aerobic energetic space when compared to the more quiescent and glycolytic status of the ciPTEC *CTNS^∆57kb^* line (Figure 4H). Furthermore, the corrected ciPTEC *CTNS*^-/-^ clones showed a reduction in glycolytic index when compared to the ciPTEC *CTNS*^-/-^ line while no improvement was found in their spare respiratory capacity (Figure 4I-J). Further analysis showed an improved glycolytic status and spare respiratory capacity of the corrected ciPTEC *CTNS^∆57kb^* derived clones, when compared to the ciPTEC *CTNS^∆57kb^* line (Figure 4K-L). Overall, these results suggest that functional restoration of *CTNS* leads to improvements in the energetic space and the mitochondrial respiratory capacity in the ciPTEC *CTNS^∆57kb^* derived clones, with a similar trend in the ciPTEC *CTNS*^-/-^.

## 4. Discussion

In this study, we aimed to restore the function of the *CTNS* gene in two cystinotic ciPTEC lines by re-introducing the wild-type promoter and first 10 exons of the gene using a CRISPR/Cas9 HITI approach. Our results showed the efficient and precise insertion of the repair templates in both cystinotic cell models, resulting in the restoration of functional *CTNS* gene, significant reduction of lysosomal cystine accumulation and improvement in mitochondrial health.

### Repair template design and insertion efficiencies

Currently most gene therapy approaches rely on the random insertion of a transgene using a viral vector. Limitations of this system are linked to variable efficiency in reaching the target tissue, insertional mutagenesis if the dose is too high, loss of transgene expression over time, adverse immune response triggered by the vector and inability of a transgene to produce multiple isoforms [35]. To overcome some of these limitations we designed a virus and plasmid-free repair system using recombinant Cas9 protein, synthetic gRNA, combined with a peptide delivery. The *CTNS* repair construct contains the first ten exons of the *CTNS* gene driven by the endogenous human promoter sequence of the *CTNS* gene. Precise integration into the genome between exon 10 and 11 allows the natural splicing of exon 11 and 12 and dynamic expression of the two main isoforms of the gene (cystinosin and cystinosisn-LKG), as well as expression of 3’UTR sequences required for mRNA transport and stability. Additionally, the HITI approach ensures that the insertion process can take place throughout the cell cycle, including non-dividing cells, in contrast to homology directed repair insertion strategies [36].

The 3.2Kb mCherry_Superexon HITI repair template showed insertion efficiencies ranging from 6% to 16% in our cystinotic cell lines, while the 1.7Kb mCherry HITI showed insertion efficiencies of 48% to 63%. Based on the clonal cell expansion, the smaller *CTNS* Superexon (1.7Kb) also resulted in high levels of integration (in 76% of clones), supporting the notion that a smaller fragment is more effective. This is most likely linked to our delivery method, which is more effective with smaller template sizes. With perfect delivery the repair efficiencies might be even higher.

After pooling all cells transfected with the 3.2Kb mCherry_Superexon HITI template and FACS sorting, we observed a rapid loss of mCherry expression over time, while the expression of *CTNS* remained constant and did not correlate with the loss of mCherry expression (Figure 3E). As both mCherry and *CTNS* are inserted in the same location in the genome, this effect is most likely caused by silencing of the CMV promoter in our cells [37–42]. These findings therefore support the notion that transgenes should be supplied with their endogenous control elements in future clinical applications of gene therapy [43].

### Phenotypical restoration of the cystinotic phenotype

Unlike other CRISPR approaches, HITI has a built-in error-correction mechanism[28] which we confirmed with >90% of our clones inserted the templates in the forward direction. Additionally, we showed that the genetic correction of one allele is sufficient to improve the phenotype, which was expected since both *CTNS* alleles need to be depleted to develop cystinosis. After the successful insertion of the HITI repair templates, we observed that an increase in *CTNS* mRNA expression as low as 5-10% was able to decrease lysosomal cystine accumulation in both ciPTEC *CTNS*^-/-^ and ciPTEC *CTNS^∆57kb^* derived clones, suggesting that the presence of a small amount of the wild-type cystinosin transporter was sufficient to restore cystinosin function. This phenomenon is in line with previous clinical and biochemical studies, which reported that patients with mild mutations that allowed for a small residual *CTNS* mRNA expression showed lower levels of cystine accumulation, either at an intermediate-cystinosis phenotype or at healthy values [44,45].

Previous studies have explored the downstream nephropathic cystinosis phenotype beyond the lysosomal cystine accumulation to provide more druggable targets for this disease [9–11,46], including mitochondrial function [47–50]. Mitochondrial respiration defects can lead to an increased oxidative stress, increased GSH levels, and a decrease in cell energy production, which has been linked to the development and exacerbation of other metabolic diseases as well, such as diabetes, obesity, cardiovascular diseases, and nephropathic cystinosis. Our findings confirm that our gene repair approach was able to restore the mitochondrial energetic space of the corrected clones, including improvements in maximal and spare respiration, proton leak and glycolytic status.

### Future perspectives

Gene repair therapies can potentially offer a unique opportunity by restoring cellular function and promoting the regenerative capacity of the kidney. While our *in vitro* gene therapy has produced promising results for cystinosis, several challenges need to be addressed before this approach can be developed into a viable gene therapy for patients.

Delivery of genes to the kidneys is notoriously difficult [51] which is why most gene therapies focus on other organs or tissues. For cystinosis, it was shown that a hematopoietic stem cell (HSC) transplantation in mice (from a wildtype mouse into a *CTNS*^-/-^ recipient) led to integration of wild type cells in various organs, effectively reducing their cystine content (by 43–93%) and preventing kidney disease [52,53]. *Ctns* kidney expression levels of 13% of the wild type levels already resulted in a full preservation of kidney function and wild type cells were also found to exchange cystinosin with neighboring cells in a process called cross-correction [54]. The homing and growth of *Ctns* wild type cells was much more efficient in cystinosin deficient tissues, suggesting that an autologous transplant of *ex vivo* repaired *CTNS*^-/-^ HSC may be able to reduce cystine levels with only mild conditioning and without irradiation or immunosuppressive therapy. Currently, a human clinical trial is ongoing to determine the safety and efficacy of a novel therapy in which *CTNS* is expressed in autologous HSC after lentiviral transduction. If successful, the next step may be to use CRISPR HITI for controlled integration in HSC.

The *ex-vivo* gene therapy approach may be a solution for cystinosis and other lysosomal storage diseases, but for other genetic (kidney) diseases direct targeting of the affected cells may be unavoidable. In this case *in vivo* gene therapy would be the preferred approach. Here, the targeting and safe delivery of genetic constructs will be the biggest challenge to be solved in the years to come but has the potential to change medicine as we know it.

## Author Contributions

Conceptualization, Elena Sendino Garví, Amer Jamalpoor, Patrick Harrison, Rosalinde Masereeuw and Manoe Janssen; Data curation, Elena Sendino Garví; Formal analysis, Elena Sendino Garví, João Faria and Suzan Thijssen; Funding acquisition, Rosalinde Masereeuw and Manoe Janssen; Investigation, Elena Sendino Garví, João Faria and Carla Pou Casellas; Methodology, Elena Sendino Garví, João Faria, Carla Pou Casellas, Suzan Thijssen, Amer Jamalpoor, Patrick Harrison and Manoe Janssen; Project administration, Rosalinde Masereeuw and Manoe Janssen; Resources, Rosalinde Masereeuw and Manoe Janssen; Supervision, Rosalinde Masereeuw and Manoe Janssen; Validation, Elena Sendino Garví, João Faria, Carla Pou Casellas and Manoe Janssen; Visualization, Elena Sendino Garví and Elise Wubbolts; Writing – original draft, Elena Sendino Garví and Elise Wubbolts; Writing – review & editing, Elena Sendino Garví, Elise Wubbolts, Amer Jamalpoor, Patrick Harrison, Rosalinde Masereeuw and Manoe Janssen.

## Institutional Review Board Statement

The cell lines used in this study were obtained from urine samples donated by human volunteers. The study was approved by the institutional (Radboudumc) ethics board. Informed consent was obtained from all human donors and the collection of these samples was approved by the committee on research involving human subjects (Commissie Mensgebonden Onderzoek) of the Radboudumc. Research was conducted in accordance with the principles embodied in the Declaration of Helsinki and in accordance with local statutory requirements. The ciPTEC line was developed in 2008 at Radboudumc from a healthy cell line is patented as ‘A novel conditionally immortalized human proximal tubule cell line expressing functional influx and efflux transporters’, Patent P6057046PCT; PCT/EP2016/080026, by Radboudumc. Since 2019, Radboudumc gave the exclusive license for the use of the cell line by Cell4Pharma, who now provides the cells; www.Cell4Pharma.nl (Nijmegen, The Netherlands, MTA #A16-0147).

## Informed Consent Statement

Not applicable.

## Acknowledgments

The authors would like to gratefully thank Georgia Avramidou for the input provided at early stages of the experimental phase. Additionally, we thank Sabbir Ahmed and Rolf Sparidans for the training and support provided for the LC-MS measurements.

## Appendix A

**Figure A1.**
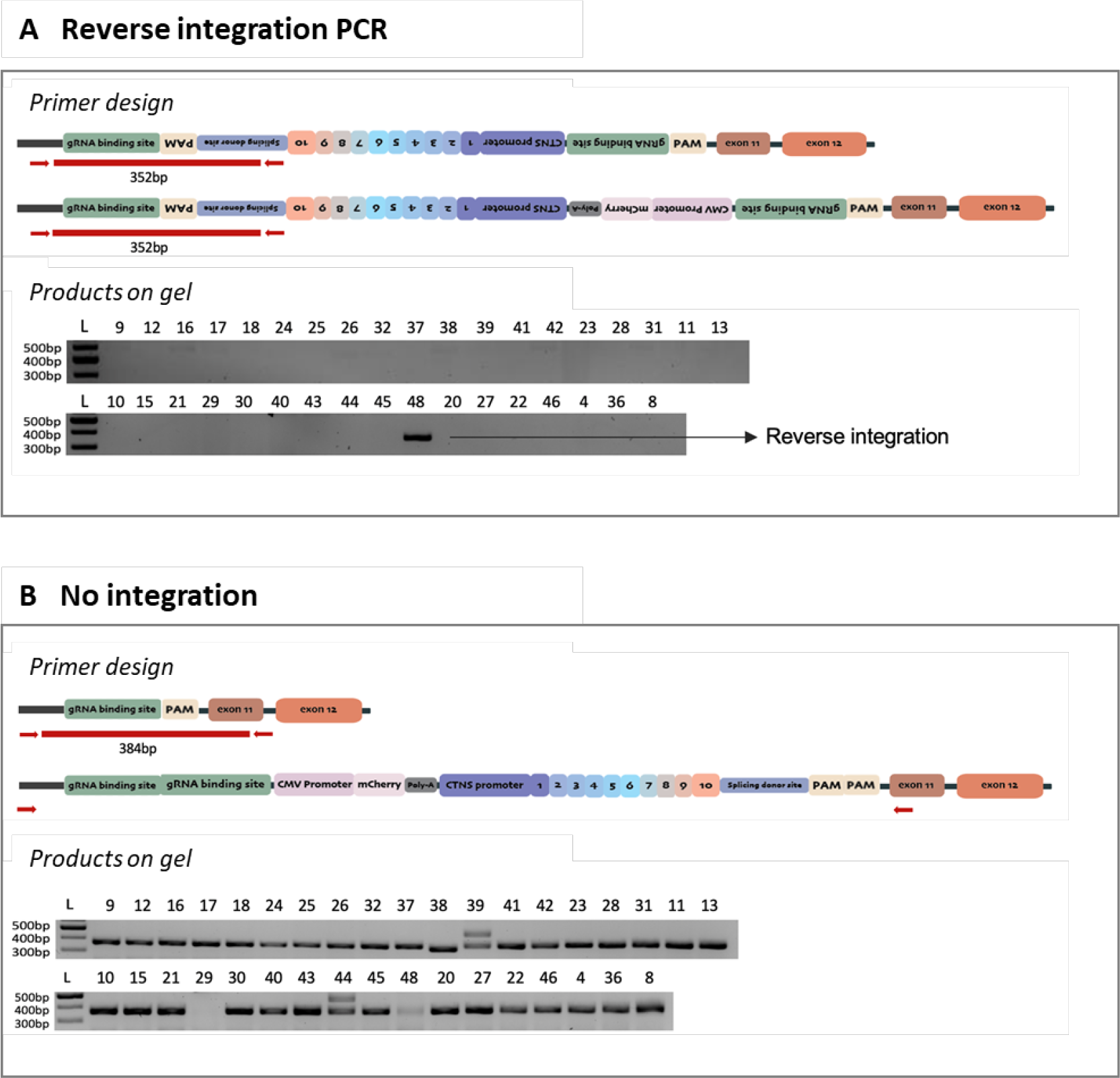
The *CTNS* repair templates show low levels of reverse integration and predominantly monoallelic integration. (**A**) Primers were designed to detect a 352bp band when the repair templates were inserted in the reversed direction. Only one clone inserted the template in the reversed direction. (**B**) Primers were designed to detect a band of 384bp when the is no integration.

